# Electrophysiological comparison of left versus right stellate ganglia neurons

**DOI:** 10.1101/2024.10.29.620825

**Authors:** Arie Verkerk, Carol Ann Remme, Molly O’Reilly

## Abstract

**Background:** The stellate ganglia of the peripheral autonomic nervous system innervate the heart and continuously fine-tune cardiac function to meet physiological demands. The right stellate ganglion (RSG) predominantly innervates the sinoatrial node and has functional effects on chronotropy/heart rate, whereas the left stellate ganglion (LSG) has predominance in the ventricular myocardium and impacts inotropy/contractility. Whilst the innervation patterns and functional consequences of block and stimulation are well-documented, basic electrophysiological characterisation and single-cell comparison of RSG and LSG neurons has not been performed. In addition, sex differences in stellate ganglion action potential (AP) parameters may exist, but remain as yet unknown.

**Methods/Results:** Here we characterise the electrical properties of enzymatically isolated mouse stellate ganglia neurons using the patch clamp technique. Using 500 ms depolarising pulses of varying amplitude, we provide detailed characterisation of basic AP properties and their correlations. We reveal that there are two populations of neurons in terms of their AP firing properties (phasic or tonic firing), with the majority (67%) firing with a phasic pattern. When all recordings were pooled, tonic neurons had a significantly larger AP amplitude (85 ± 3.0 vs 76 ± 2.4 mV) and overshoot (28 ± 1.8 vs 19 ± 1.8 mV) compared to phasic neurons (P<0.05). Moreover, phasic neurons did not fire spontaneously, whereas 50% of tonic neurons did, and more often presented with anodal break excitation (P<0.05). When male vs female neurons were compared, males had a more negative minimum diastolic potential (MDP; -55 ± 1.7 vs -47 ± 3.0 mV, P<0.05). When LSG vs RSG neurons were compared, the RSG had a more negative resting membrane potential (V_rest_; -60 ± 1.5 vs -54 ± 1.3 mV, P<0.05). All other AP parameters did not differ significantly between these groups.

**Conclusions:** The RSG and LSG contain a similar proportion of phasic and tonic firing neurons. A significant difference was observed in the V_rest_ of RSG vs LSG neurons, and in the MDP of male vs female neurons. However, all other AP parameters were similar. This suggests that the LSG and RSG can be combined irrespective of sex when investigating the electrophysiological properties of these distinct anatomical structures in healthy and disease conditions.

## Introduction

The stellate ganglia are vitally important anatomical structures in respect to cardiac function. They are part of the peripheral autonomic nervous system, located alongside the spinal cord, and contain the vast majority of the sympathetic neurons that project directly to the heart. During any instances of sympathetic activity (exercise, emotion, or fight-or-flight response) the neurons of the stellate ganglia, and their cardiac-projecting neurites, become activated. This fine-tunes the functioning of the heart, enhancing sympathetic tone, modifying neurotransmitter release, and increasing cardiac output to meet the demands of the new physiological state.

In numerous cardiac diseases, the stellate ganglia are recognised as a pathophysiological contributor and thus a target for treatment. In inherited arrhythmia syndromes such as Catecholaminergic Polymorphic Ventricular Tachycardia (CPVT) and Long-QT Syndrome (LQTS), surgical removal of the stellate ganglia (stellectomy) is a known treatment approach that yields positive outcomes (Schwartz et al, 2004; Wilde et al, 2008; Schwartz and Ackerman, 2022). Moreover, recent fundamental studies reveal there to be functional alterations of the stellate ganglia in pro- (Larsen et al., 2016) and pre-hypertensive (Davis et al, 2020) animal models.

The right and left stellate ganglia have overlapping but also distinct cardiac innervation patterns. The right stellate ganglion (RSG) has predominant innervation of the pace-setting sinoatrial node of the heart, whereas the left stellate ganglion (LSG) predominantly innervates the ventricular myocardium. Consequently, the ganglia have differing functional effects in terms of their primary physiological impact on chronotropy/heart rate (RSG) or inotropy/contractility (LSG) (Yanowitz et al, 1966; Zandstra et al, 2021; Li 2022).

Despite recognition of their differing physiological roles, few studies have investigated the LSG and RSG in detail, and often in studies of disease models neurons from the LSG and RSG are pooled. Whilst the innervation patterns and functional consequences of block and stimulation are well-documented, as well as their neurochemical profiles (Bayles et al, 2018), basic electrophysiological characterisation and comparison of single cell RSG and LSG neurons is currently lacking. Moreover, a transcriptomic study of the LSG has demonstrated sex differences in the expression of genes that encode ion channels (Bayles et al, 2018). However, any resulting electrical differences have as yet not been investigated. This study aims to fill this knowledge gap by providing action potential (AP) characterisation and comparison of the LSG and RSG in both male and female mice - investigating whether electrical differences exist and if it is appropriate to pool the two distinct anatomical structures when performing comparative disease studies.

## Methods

### Animals

Wild-type C57Bl6j mice were used for the experiments detailed herein - male and female, 2-6 months. Housing, handling, and experiments were performed in agreement with the Institutional guidelines.

### Stellate ganglion neuron isolation

Following terminal anaesthesia (4% isoflurane inhalation in O_2_), RSG and LSG were separately removed and placed in ice-cold PBS. Single RSG and LSG neurons were isolated by an enzymatic dissociation procedure. Ganglia were transferred to a nominally Ca^2+^-free Tyrode’s solution (20°C) (pH 7.4; NaOH) containing (in mM): NaCl 140, KCl 5.4, CaCl_2_ 0.01, MgCl_2_ 1, glucose 5.5, HEPES 5 as well as Liberase TM (26 U/ml) and Elastase (211 U/ml) enzymes. The tissue was gently agitated in a shaking water bath at 37°C for 28 minutes. Subsequently, ganglia were washed in nominally Ca^2+^-free Tyrode’s solution (20°C), and thereafter in normal Tyrode’s solution (20°C) (pH 7.4; NaOH) containing (in mM): NaCl 140, KCl 5.4, CaCl_2_ 1.8, MgCl_2_ 1, glucose 5.5, HEPES 5. Finally, ganglia were transferred to B-27 Plus Neuronal Culture System (Gibco) media (20°C) and single cells were obtained by manual trituration using fire-polished glass pipettes. Single cells were plated on coverslips coated with 100 ug/ml poly-d-lysine and 10 ug/ml laminin. Neurons were left to adhere in a 37°C incubator overnight before experiments were performed the subsequent day, and such a short culturing period does not affect the electrical phenotype of stellate ganglia neurons (Davis et al., 2020). 2 mice were used for each isolation. Measurements were only included in analysis when recordings were obtained from both the left and right stellate from the same isolation. A total of 6 different isolations were included in the analysis.

### Patch clamp electrophysiology

Coverslips were transferred to a recording chamber and were continually superfused with normal Tyrode’s solution (37°C). APs were recorded using the amphotericin perforated patch clamp technique with an Axopatch 200B amplifier (Molecular Devices, Sunnyvale, CA, USA). Data acquisition was realized using a CED micro1401 driven by custom-made acquisition software (Axograph) and data analysis was performed with custom software (MacDaq, version 15.4; kindly provided by Antoni C. G. van Ginneken). Signals were low pass filtered with a cut-off frequency of 5 kHz and digitized at 50 kHz. Patch pipettes were pulled from borosilicate glass (TW100F-3, World Precision Instruments Germany Gmb) using a vertical microelectrode puller (PC-100; Narishige Scientific Instrument, Japan) and had tip resistances of 1.5–3 MΩ after filling with the pipette solution as indicated below. All potentials were corrected for the estimated liquid junction potential (Barry and Lynch, 1991).

Patch pipettes were filled with a solution containing (in mM): K-gluconate 125, KCl 20, NaCl 5, amphotericin-B 0.44, HEPES 10, pH 7.3 (KOH). APs were evoked by 500 ms depolarising current pulses of varying amplitude (0–200 pA, in 50 pA steps). We counted the number of APs during a 500 ms stimulus. Further, we analysed AP parameters as described previously (Ten Hoope et al., 2018) and cells which did not overshoot the zero potential level were excluded. The resting membrane potential (V_rest_) was defined as the potential immediately before the depolarising pulse. From the first AP of the 150 pA depolarising pulse, we measured the: AP overshoot, AP amplitude (APA) as the difference between V_rest_ and overshoot, maximal AP upstroke velocity (V_max,dep_), maximal AP repolarisation velocity (V_max, rep_), AP duration (APD) at 50% repolarisation (APD_50_) and the minimum diastolic potential (MDP) as the most negative hyperpolarisation voltage following the first AP. Finally, we tested anodal break excitation, i.e. AP generation after a hyperpolarising pulse, by a 100 pA, 500 ms hyperpolarising pulse. All analysed AP parameters are schematically indicated in Figure 1A.

**Figure 1.**
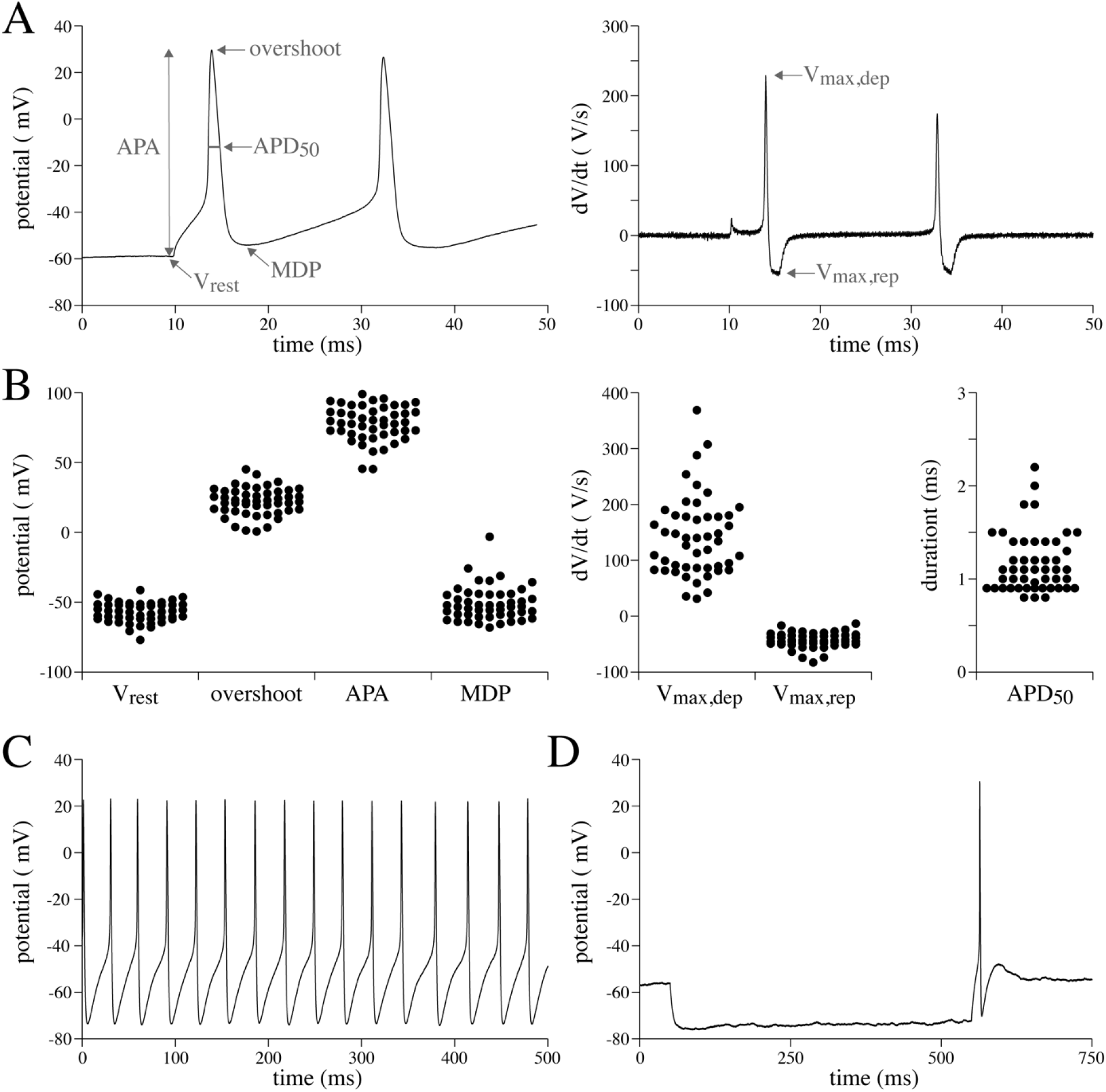
Basic action potential (AP) characteristics of stellate ganglion neurons. A, Example of APs evoked by a 150 pA depolarising pulse (left panel) and the first derivative of these APs (right panel). The analysed parameters are schematically indicated. B, Dot plots of all AP parameters of 48 neurons. C, Typical example of spontaneous APs (without current injection). D, Typical example of anodal break excitation in response of a 100 pA hyperpolarising pulse. V_rest_=potential 5 ms before the first AP, APA=AP amplitude, MDP=maximal diastolic potential after the first AP, V_max,dep_=maximal AP upstroke velocity, V_max,rep_=maximal velocity of AP repolarisation, APD_50_=AP duration at 50% of repolarisation.

### Statistics

Statistical analysis was carried out with SigmaStat 3.5 (Systat Inc, St. Louis, MO). Normality and equal variance assumptions were tested with the Kolmogorov-Smirnov and Levene median tests, respectively. Two groups were compared using the Fisher exact test, the unpaired t-test or, if normality and/or equal variance tests failed, the Mann-Whitney rank-sum test. One-way and Two-way repeated measures (RM) ANOVA, followed by the Students-Newman-Keuls post hoc test was used to test the significance for the data in Figure 3C. Linear relationships were analysed using the Pearson correlation coefficient (R) and significance level. Weak, moderate and strong relationships were defined R values of <0.3, 0.3-0.5, and 0.5>, respectively. *P*<0.05 was considered statistically significant.

## Results

### Basic action potential properties of stellate ganglion neurons

#### Overview of AP parameters

In Figure 1, we summarized the general AP properties of all measured stellate ganglion neurons. For this purpose, all recordings obtained from left and right stellate ganglion neurons of both male and female mice were pooled. Figure 1A, left panel, shows typical AP recordings evoked by a 150 pA, 500 ms long depolarising pulse, i.e., the pulse where all measured cells evoked at least one AP. Figure 1A, right panel, shows the first derivative of the AP, reflecting the change in voltage per second (dV/dt). Analysed AP parameters are schematically indicated. Figure 1B depicts dot plots of the analysed AP properties of the first evoked AP. V_rest_ was between -40 and -80 mV and overshoot ranged between 0 and 50 mV. Consequently, APA was typically 50 to 100 mV. V_max,dep_ was quite variable with values between 30 and 400 V/s, indicating that the main inward current underlying the AP upstroke, i.e., the sodium current, may differ largely between neurons. V_max,rep_, due to outwardly directed potassium currents, was between -30 and -70 V/s. The APD_50_ was typically short and ranged between 0.8 and 2.5 ms. MDP was slightly more depolarised than the V_rest_. Without current injection, we found that 7 out of 48 neurons (14.6%) showed spontaneous AP generation, with typical examples shown in Figure 1C. Moreover, anodal break excitation in response to a 100 pA hyperpolarisation pulse was observed in 20 out of 48 neurons (41.7%; Figure 1D).

#### Relationships between AP parameters

It is evident from Figure 1B that there is some variation in AP parameters between neurons. Figure 2A shows typical AP recordings from two separate neurons obtained from the same right stellate ganglion isolation, displaying a clear difference in V_rest_ between both cells. The AP with the most negative V_rest_ had the fastest AP upstroke velocity, but also the fastest repolarisation and shortest AP. In addition, the MDP in this cell was more negative compared to the other depicted neuron. Next, we determined if this is a consistent finding by testing for relationships between AP parameters. Therefore, we plotted linear fits and performed analysis of the Pearson correlation coefficients (R) and significance (Figure 2B-G). All R’s between the AP properties are summarized in Figure 2G, and it is evident that there are multiple strong and moderated linear associations between the variables. For example, the V_rest_ shows a linear relationship with the AP amplitude (APA) with an R of -0.70 (P<0.001) (Figure 2B,G), thus indicating a strong negative correlation. This may be partially related to our definition of APA, since we defined it as the difference between V_rest_ and overshoot. More importantly, V_rest_ also shows linear correlations (P<0.05) with V_max,dep_ (Figure 2C,G), V_max,rep_ (Figure 2D,G), APD_50_ (Figure 2E,G) and MDP (Figure 2F,G), although these can be considered as moderate. These correlations are likely caused by neuronal membrane current properties. V_rest_ affects importantly the sodium current responsible for the AP upstroke, and a more depolarised V_rest_ will result in less available sodium channels since many channels will be in the inactivated state. As a result, V_max,dep_ will be smaller. Similarly, a more depolarised V_rest_ will reduce the availability of the I_A_ current, i.e., the transient outward potassium current, which is an important repolarising current in neurons. Consequently, the V_max,rep_ becomes smaller resulting in longer APs with more positive MDPs, as indicated by the strong correlations of these parameters. There is also a strong negative correlation between V_max,dep_ and V_max,rep_ (Figure 2G), but as mentioned above this is likely due to their underlying channel properties and the common relation with V_rest_.

**Figure 2.**
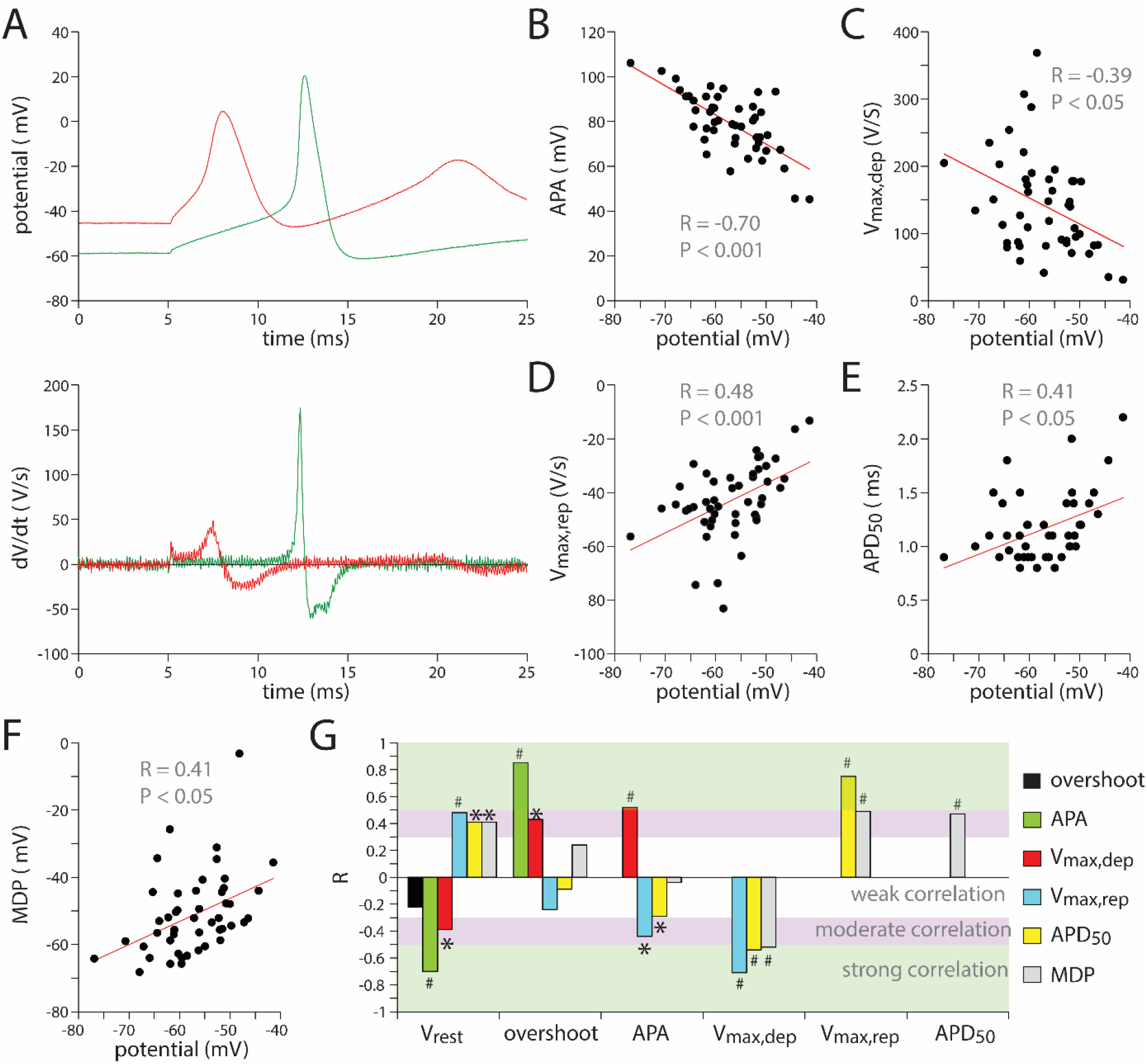
Correlations between AP parameters. A, Two examples of APs with different V_rest_ evoked by a 150 pA depolarising pulse (top panel) and the first derivative of these APs (bottom panel). B to F, Linear relationships (red solid lines) of V_rest_ with APA (B), V_max,dep_ (C), V_max,rep_ (D), APD_50_ (E), and MDP (F). R indicates the Pearson correlation coefficient. G, R values of all linear relationships between AP parameters. Green, pink and white horizontal bars indicate the strength of the correlations. *P<0.05, ^#^P<0.001.

#### AP firing patterns

In the sections above, we focused on the first AP evoked by the 150 pA depolarising pulse. However, looking into the firing pattern during the complete 150 pA depolarising pulse in more detail revealed two distinct populations of neurons, as shown in the typical AP examples of Figure 3A. The majority of the neurons (32 out of 48) fired with a ‘phasic’ pattern – firing a small number of APs (<4, Figure 3B) before becoming quiescent for the rest of the 150 pA, 500 ms depolarising pulse. The remaining 33% of the neurons fired with a ‘tonic’ pattern – firing almost continuously throughout the same depolarising stimulus and the number of APs during the 150 pA, 500 ms depolarising pulse is typically >9 (Figure 3B). The firing frequencies of both the phasic and tonic neurons significantly increased in response to increasing depolarising pulses (P<0.05, One-way RM ANOVA), but that of phasic neurons remained relatively low at all depolarising pulses (Figure 3C). Consequently, phasic and tonic firing patterns differ significantly (P<0.05, Two-way RM ANOVA) at all tested depolarising pulses (Figure 3C). Interestingly, phasic neurons did not fire spontaneously, whereas almost 50% of tonic neurons did fire spontaneously without any current injection (7 out of 16 cells) (Figure 3D). The number of cells with anodal break excitations was also significantly higher in tonic neurons (Figure 3D).

When analysing the parameters of the first AP at the 150 pA depolarising pulses, tonic neurons were found to have a significantly larger overshoot and APA compared to phasic neurons (Figure 3E). All other AP parameters (V_rest_, MDP, V_max,dep_, V_max,rep_ and APD_50_) did not differ significantly between phasic and tonic neurons.

**Figure 3.**
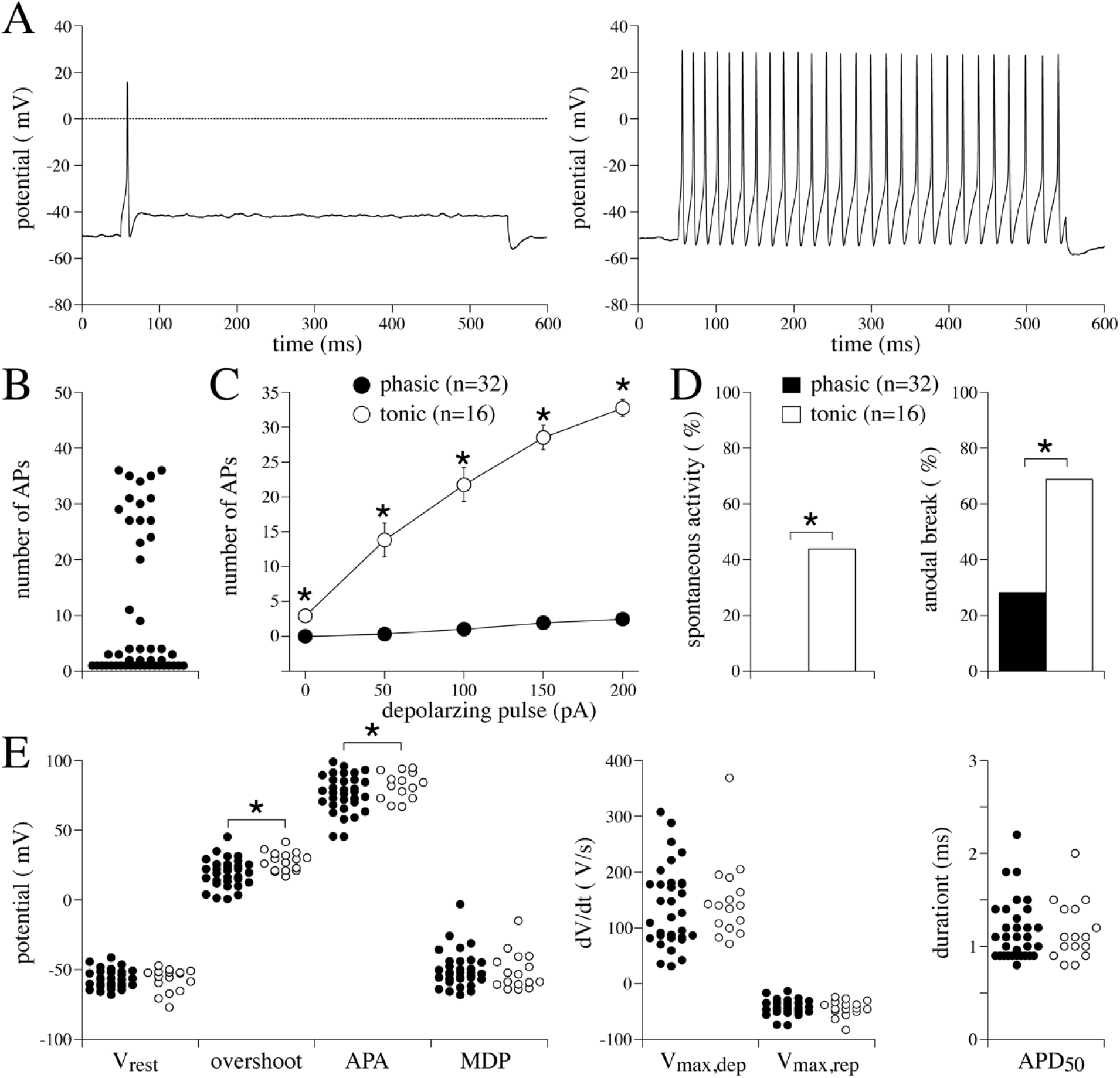
AP parameters of phasic and tonic neurons. A, Two examples of firing pattern evoked by a 150 pA, 500 ms depolarising pulse from one left stellate ganglia isolation. B, Dot plot of the number of APs during the 150 pA, 500 ms depolarising pulse. C, Number of APs vs the depolarising pulses for phasic and tonic neurons *P<0.05 (Two-way RM ANOVA). D, Percentages of phasic and tonic neurons with spontaneous APs and anodal break excitation *P<0.05 (Fisher exact test). E, Dot plots of the AP parameters in phasic and tonic neurons. *P<0.05 (unpaired t-test).

### AP properties of male versus female stellate ganglion neurons

In the above section, we pooled the AP data of female and male mice. However, if sex differences in stellate ganglion neurons exist, this approach might be questionable. Hence, we next compared the basic firing patterns and AP properties of male and female mice. As shown in Figure 4A, no significant differences in the number of APs and ratio of phasic neurons were observed between the two sexes. In addition, neither the number of cells with spontaneous activity nor that with anodal breaks differed (Figure 4B). Neurons of male mice have a slightly more negative MDP (*P*<0.05), but all other parameters are not affected by sex (Figure 4C).

**Figure 4.**
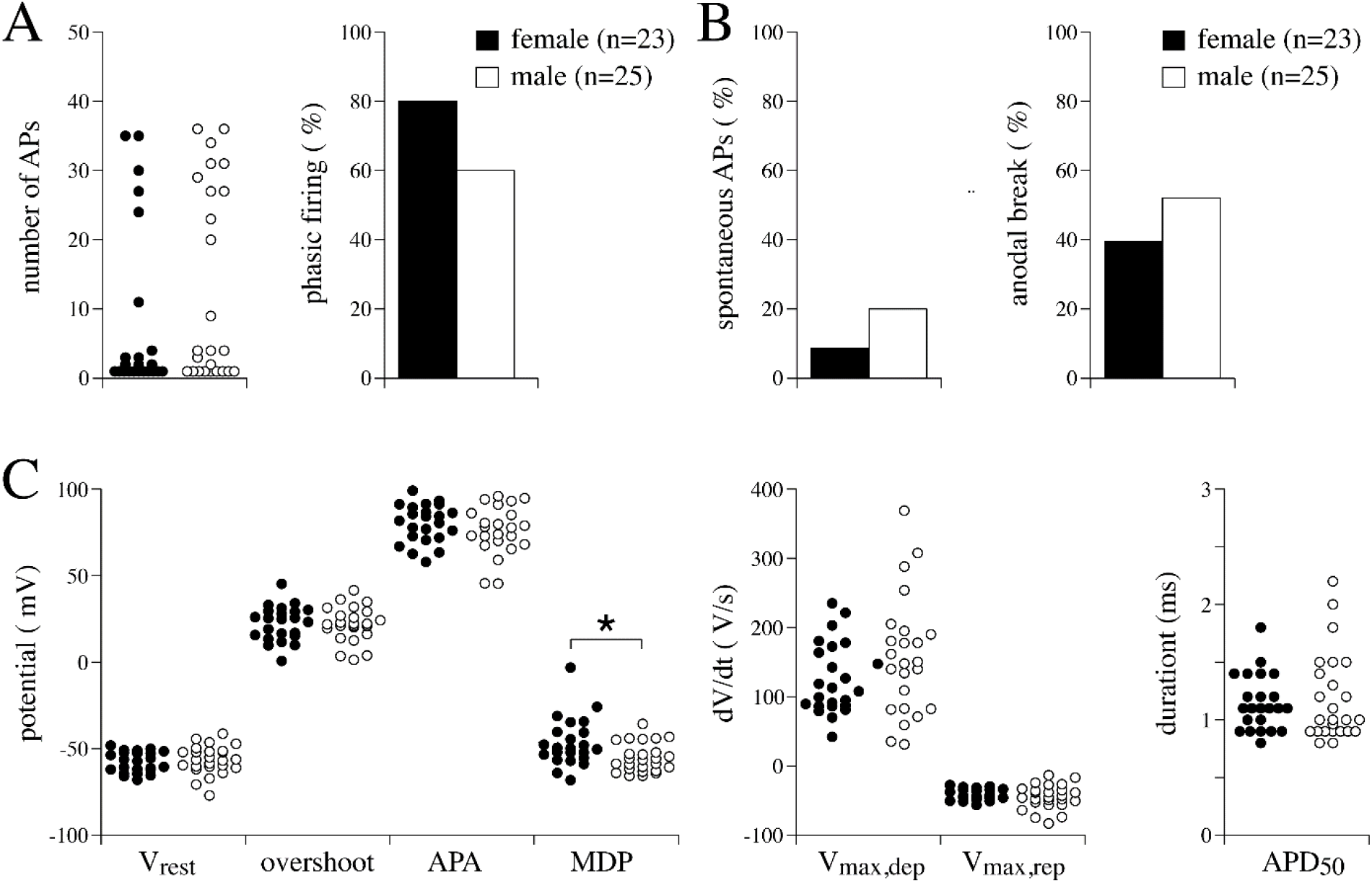
AP parameters of female and male neurons. A, number of APs (left panel) and percentage of phasic firing neurons (right panel) during the 150 pA depolarising pulse of female and male mice. B, Percentage of neurons with spontaneous APs (left panel) and anodal break excitation (right panel) of female and male mice. C, Dot plots of the AP parameters in female and male neurons. *P<0.05 (Mann-Whitney Rank sum test).

### AP properties of left versus right stellate ganglion neurons

Finally, we compared the firing pattern and AP properties of isolated neurons from the LSG and RSG, pooling male and female recordings. Neither the number of APs nor the ratio of phasic neurons differs between the RSG and LSG. In addition, the number of cells with spontaneous AP generation and anodal break excitation was similar (Figure 5B). RSG neurons have a slightly more negative V_rest_ (P<0.05), but all other AP properties were not significantly different (Figure 5C).

**Figure 5.**
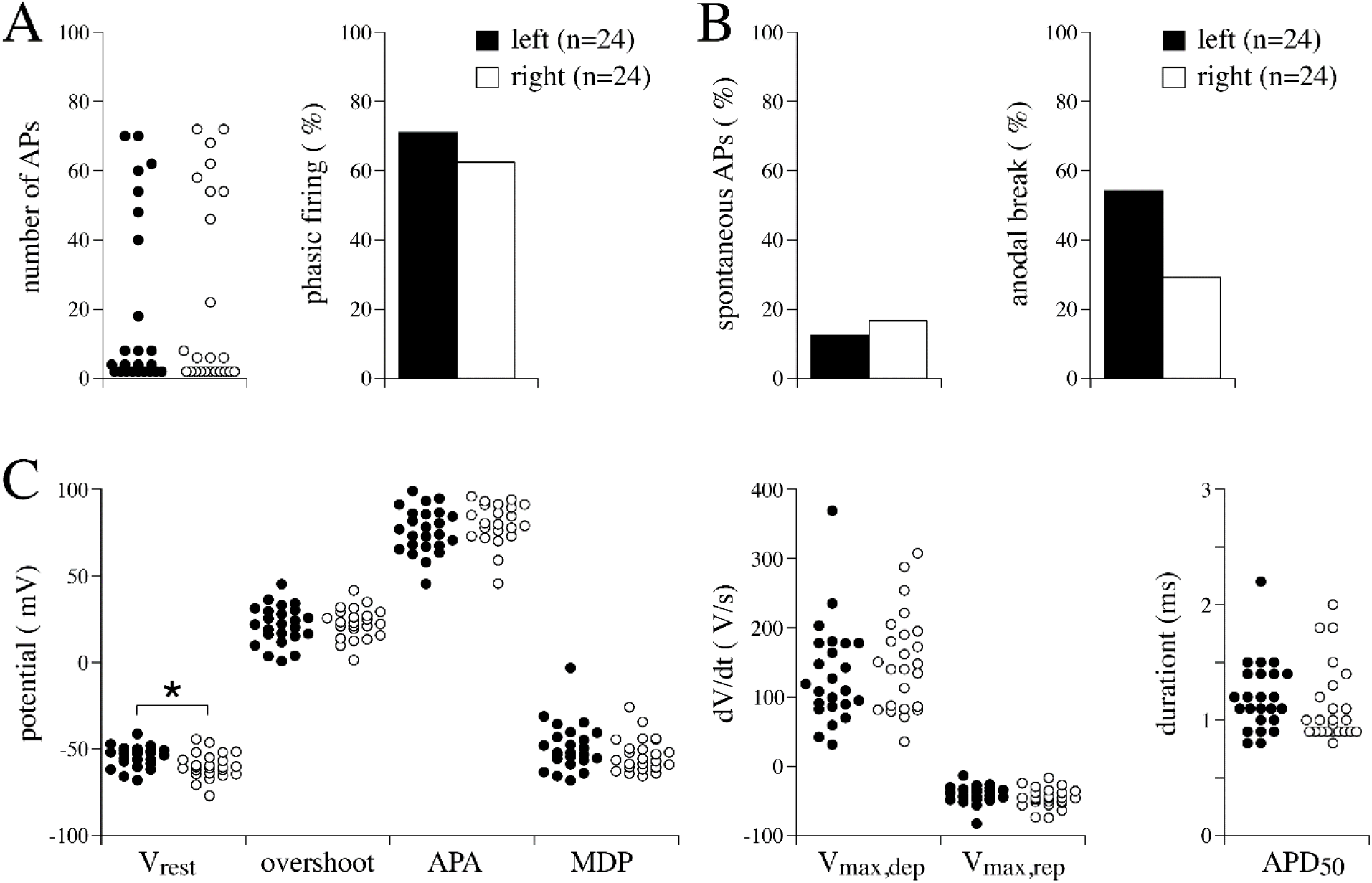
AP parameters of left and right ganglia neurons. A, number of APs (left panel) and percentage of phasic firing neurons (right panel) during the 150 pA depolarising pulse in left and right ganglion neurons. B, Percentage of neurons with spontaneous APs (left panel) and anodal break excitation (right panel) of left and right neurons. C, Dot plots of the AP parameters. *P<0.05 (t-test).

## Discussion

Here we document for the first time the comparative basic electrophysiological profiles of isolated neurons from the left (LSG) versus the right stellate ganglion (RSG) of mice. Detailed analysis of basic action potential (AP) parameters (V_rest_, V_max,dep_, V_max,rep_, APA, overshoot, MDP, and APD_50_) revealed a number of relationships and correlations, as expected from the underlying ionic basis of these measures. When pooling the data of all neurons recorded from the LSG and RSG we revealed two distinct neuronal populations based on AP firing patterns, namely tonic or phasic type neurons, as has been reported previously in the stellate ganglia of rats (Davis et al., 2020), and in the sympathetic chain of guinea pigs (Cassell et al., 1986). We also reveal novel differences in these neuronal subtypes including a heightened propensity for spontaneous AP firing and anodal break excitation in tonic neurons compared to phasic, and significant differences in AP amplitude and overshoot properties. It is known that these neuronal subtypes have different target tissue innervation patterns, in that tonic neurons innervate fewer postsynaptic targets than phasic (Aponte-Santiago et al, 2020); these novel electrophysiological characterisations may be important for these innervation differences.

When comparing the basic AP properties of male versus female stellate ganglia neurons, we observed a small difference in the MDP, but not in any other AP parameter. This suggests that the electrical profile of these neurons is not affected by sex, and thus the two sexes can be combined when performing comparative disease studies (O’Reilly et al., 2024). Interestingly, researchers have performed RNAseq analysis of the left stellate ganglia and identified gene expression differences between male and female C57Bl6j mice, including differences in the expression of genes that are important for encoding ion channels, such as *Kcna2* (K_v_1.2) (Bayles et al., 2018). However, here we show that these ion channel gene expression differences do not translate to differences in the baseline resting membrane potential or the excitability of these cardiac sympathetic neurons, as the researchers had speculated.

Lastly, we also compared the basic AP parameters of the LSG and RSG. Whilst most parameters were comparable, we found that RSG neurons have a significantly more negative resting membrane potential (V_rest_) than LSG neurons (−60 vs -54 mV). This subtle electrophysiological difference may pertain to the different functional roles of the two structures, with the RSG predominantly innervating and affecting the sinoatrial node and heart rate, and the LSG having predominance of ventricular myocardium innervation and contractility. Speculatively, the more negative V_rest_ in the RSG would allow for faster and more efficient triggering of APs following sympathetic activation, to rapidly induce heart rate changes upon changes in physiological demands.

## Conclusion

Here we show that the electrical profile of neurons of the LSG and RSG of males and females is similar, and thus they can be pooled in future studies. We provide a protocol for the isolation and investigation of mouse stellate ganglia neurons, which will be useful for studies of disease conditions involving genetic factors and evidence of stellate involvement. For example, in inherited arrhythmia conditions such as Catecholaminergic Polymorphic Ventricular Tachycardia (CPVT) and Long-QT Syndrome (LQTS), where stellectomy is a viable treatment option.

## Limitations

The underlying ionic basis of the observed AP differences could not be investigated, as one cannot first identify whether a neuron is phasic or tonic (current-clamp configuration) before recording their ionic currents (voltage-clamp configuration). Selective fluorescence labelling of the different types of neurons, if possible, could aid future studies to investigate this further.

## Author contributions

Conceptualization, M.O., A.O.V. and C.A.R.; Data collection, M.O.; Data analysis, M.O. and A.O.V.; Funding acquisition, M.O. and C.A.R.; Visualization, M.O. and A.O.V; Writing, M.O., A.O.V. and C.A.R. All authors have read and agreed to the published version of the manuscript.

## References

Aponte-Santiago NA, Ormerod KG, Akbergenova Y, Littleton JT. Synaptic Plasticity Induced by Differential Manipulation of Tonic and Phasic Motoneurons in Drosophila. J Neurosci. 2020 Aug 12;40(33):6270–6288. doi: 10.1523/JNEUROSCI.0925-20.2020. Epub 2020 Jul 6. PMID: 32631939; PMCID: PMC7424871.

Barry, PH and Lynch, JW. Liquid junction potentials and small cell effects in patch-clamp analysis. J. Membr. Biol. 1991, 121, 101–117. doi: 10.1007/BF01870526.

Bayles RG, Olivas A, Denfeld Q, Woodward WR, Fei SS, Gao L, Habecker BA. Transcriptomic and neurochemical analysis of the stellate ganglia in mice highlights sex differences. Sci Rep. 2018 Jun 12;8(1):8963. doi: 10.1038/s41598-018-27306-3. Erratum in: Sci Rep. 2019 Jun 26;9(1):9506. doi: 10.1038/s41598-019-45211-1. PMID: 29895973; PMCID: PMC5997635.

Cassell JF, Clark AL, McLachlan EM. Characteristics of phasic and tonic sympathetic ganglion cells of the guinea-pig. J Physiol. 1986 Mar;372:457–83. doi: 10.1113/jphysiol.1986.sp016020. PMID: 2425087; PMCID: PMC1192774.

Davis H, Herring N, Paterson DJ. Downregulation of M Current Is Coupled to Membrane Excitability in Sympathetic Neurons Before the Onset of Hypertension. Hypertension. 2020 Dec;76(6):1915–1923. doi: 10.1161/HYPERTENSIONAHA.120.15922. Epub 2020 Oct 12. PMID: 33040619; PMCID: PMC8360673.

Larsen HE, Lefkimmiatis K, Paterson DJ. Sympathetic neurons are a powerful driver of myocyte function in cardiovascular disease. Sci Rep. 2016 Dec 14;6:38898. doi: 10.1038/srep38898. PMID: 27966588; PMCID: PMC5155272.

Li YL. Stellate Ganglia and Cardiac Sympathetic Overactivation in Heart Failure. Int J Mol Sci. 2022 Nov 1;23(21):13311. doi: 10.3390/ijms232113311. PMID: 36362099; PMCID: PMC9653702.

O’Reilly M, De Waal T, Casini S, Remme CA. Ventricular hyperinnervation in a mouse model of catecholaminergic polymorphic ventricular tachycardia, Cardiovascular Research, Volume 120, Issue Supplement_1, May 2024, cvae088.098, 10.1093/cvr/cvae088.098

Schwartz PJ, Priori SG, Cerrone M, Spazzolini C, Odero A, Napolitano C, Bloise R, De Ferrari GM, Klersy C, Moss AJ, Zareba W, Robinson JL, Hall WJ, Brink PA, Toivonen L, Epstein AE, Li C, Hu D. Left cardiac sympathetic denervation in the management of high-risk patients affected by the long-QT syndrome. Circulation. 2004 Apr 20;109(15):1826–33. doi: 10.1161/01.CIR.0000125523.14403.1E. Epub 2004 Mar 29. PMID: 15051644.

Schwartz PJ, Ackerman MJ. Cardiac sympathetic denervation in the prevention of genetically mediated life-threatening ventricular arrhythmias. Eur Heart J. 2022 Jun 6;43(22):2096–2102. doi: 10.1093/eurheartj/ehac134.

Ten Hoope W, Hollmann MW, de Bruin K, Verberne HJ, Verkerk AO, Tan HL, Verhamme C, Horn J, Rigaud M, Picardi S, Lirk P. Pharmacodynamics and Pharmacokinetics of Lidocaine in a Rodent Model of Diabetic Neuropathy. Anesthesiology. 2018 Mar;128(3):609–619. doi: 10.1097/ALN.0000000000002035. PMID: 29251644.

Wilde AA, Bhuiyan ZA, Crotti L, Facchini M, De Ferrari GM, Paul T, Ferrandi C, Koolbergen DR, Odero A, Schwartz PJ. Left cardiac sympathetic denervation for catecholaminergic polymorphic ventricular tachycardia. N Engl J Med. 2008 May 8;358(19):2024–9. doi: 10.1056/NEJMoa0708006. PMID: 18463378.

Yanowitz F, Preston JB, Abildskov JA. Functional distribution of right and left stellate innervation to the ventricles. Production of neurogenic electrocardiographic changes by unilateral alteration of sympathetic tone. Circ Res. 1966 Apr;18(4):416–28. doi: 10.1161/01.res.18.4.416. PMID: 4952701.

Zandstra TE, Notenboom RGE, Wink J, Kiès P, Vliegen HW, Egorova AD, Schalij MJ, De Ruiter MC, Jongbloed MRM. Asymmetry and Heterogeneity: Part and Parcel in Cardiac Autonomic Innervation and Function. Front Physiol. 2021 Sep 16;12:665298. doi: 10.3389/fphys.2021.665298. PMID: 34603069; PMCID: PMC8481575.

